# Long-term fertilization modifies the composition of slow-growing oligotrophs more than fast-growing copiotrophs in a nutrient poor coastal plain wetland

**DOI:** 10.1101/2022.11.28.518233

**Authors:** Allison Walker, Daniya Stephens, Aied Garcia, Ariane L. Peralta

## Abstract

Wetland ecosystems are known for their carbon storage potential due to slow decomposition rates and high carbon fixation rates. However, nutrient addition from human activities affects this carbon storage capacity as the balance of fixed and respired carbon shifts due to plant and microbial communities. Ongoing atmospheric deposition of nutrients could be changing wetland plant-microbe interactions in ways that tip the balance from carbon storage to loss. Therefore, examining microbial community patterns in response to nutrient enrichment is important to understanding nutrient effects on carbon storage potential. In this study, we hypothesized that fertilization of a low nutrient ecosystem leads to increased organic carbon input from plant biomass into the soil and results in increased soil bacterial diversity and modifications to soil bacterial community composition. As such, increased soil nutrients and carbon resources provide more energy to support increased microbial growth rates, which can result in wetland carbon losses. To test this hypothesis, we used bacterial community-level and soil chemical data from the long-term wetland ecology experiment at East Carolina University’s West Research Campus (established in 2003). Specifically, we examined the extent that long-term effects of nutrient enrichment affect wetland microbial communities and plant biomass, which are factors that can affect carbon storage. We collected soil cores from fertilized and unfertilized test plots. We extracted genomic DNA from soil samples and conducted 16S rRNA targeted amplicon sequencing to characterize the bacterial community composition. In addition, we measured plant above and belowground biomass and soil carbon content. Results revealed an increase in aboveground plant biomass, soil carbon, and bacterial diversity. In contrast, belowground plant biomass and microbial biomass were similar in fertilized and unfertilized plots. To further examine bacterial community changes to nutrient enrichment, we compared the relative abundance of fast growing copiotrophic and slow growing oligotrophic bacteria of a subset of taxa putatively identified as belonging to either life history strategy. These taxa-level results revealed a decrease in oligotroph relative abundance and little to no change in copiotroph relative abundance of a subset of bacterial taxa. If there is a community-wide shift in the proportion of oligotroph to copiotroph life history strategies, this would have a negative impact on organic carbon storage since oligotrophic bacteria respire less carbon than copiotrophic bacteria over the same amount of time. Taken together, this study provided evidence that long-term nutrient enrichment influences wetland soils in ways that decrease their carbon storage potential of important carbon sinks.

## INTRODUCTION

Ecosystem functions such as nutrient and carbon cycling can benefit society by providing water and air quality improvement and climate change mitigation. But human activities such as land use change (e.g., agriculture, urbanization, deforestation) and industrialization (i.e., fossil fuel combustion, contaminant production) are causing unprecedented changes to the environment (Blaser et al. 2016). These activities add an excess of nutrients that inevitably impair important ecosystem functions by changing carbon and nutrient cycles (Davidson et al. 2012). The effects of nutrient enrichment on soils can be clearly seen in agriculturally dominated areas of the United States. Not only is fertilizer applied directly to agricultural lands, but nitrogen and other nutrients can deposit onto terrestrial and aquatic ecosystems from the atmosphere and indirectly fertilize soils. In particular, US midwestern states and eastern North Carolina represent hotspots of atmospheric nitrogen deposition (Schwede et al. 2014). Past studies reveal that nutrient enrichment can increase plant biomass carbon (Borer et al. 2017, Harpole et al. 2016) which in turn increases microbial respiration to release more carbon dioxide into the atmosphere (Koceja et al. 2021, Hoosbeek et al. 2004, Kuzyakov 2010). Therefore, understanding the extent that this nutrient addition modifies the balance of carbon fixed by plants and carbon loss through decomposition by microorganisms is critical for accurately predicting an ecosystem’s carbon storage capacity (Hoosbeek et al. 2004).

While carbon accrual into the soil from increased plant productivity is beneficial, climate change effects such as elevated atmospheric carbon dioxide and temperatures can also disrupt the carbon cycle by modifying the rate of carbon loss to the atmosphere (Borer et al. 2017, Lange et al. 2015). Previous studies have shown that high levels of carbon dioxide in the atmosphere increase plant growth leading to higher amounts of respired carbon from microbial decomposition (Jiang et al. 2020). Increased temperatures also promote higher rates of decomposition (Cavicchioli et al. 2019).

The increase of nutrients due to climate change influences the microbial community composition. For example, long term enrichment of grasslands has shown that increasing a limiting resource, such as nitrogen, decreases species diversity and increases carbon accrual (Harpole et al. 2016, Fornara and Tilman 2012). This loss of diversity can be detrimental to the environment since diversity within an ecosystem promotes stability and productivity (Borer et al. 2017). Changes to plant species diversity can differentially affect functional groups of soil microbial communities whose activities are critical for carbon storage or loss through decomposition.

The life history strategy of the soil microbes can determine the effects a changing community composition would have on the ecosystem. The two life history categories of bacteria that are relevant to this study are oligotrophic and copiotrophic as described in previous literature (Fierer et al. 2007). Because of the slow-growing nature of oligotrophic bacteria, less carbon is respired over time compared to fast-growing copiotrophic bacteria leading to greater carbon storage in soils where they are predominant (Fierer et al. 2007). In nutrient limiting environments, oligotrophic bacteria have greater growth efficiency compared to copiotrophs and outcompete them. However, excess nutrients in soil can result in the copiotrophic bacteria to grow more quickly which can eventually shift patterns of bacterial communities and ultimately influence the types and rates of microbial metabolisms (Roller et al. 2015).

Nutrient feedback on carbon cycling is especially important to examine in ecosystems that are considered effective carbon sinks. For example, wetlands take up only 7% of the surface of the earth but store 30% of the supply of carbon (USDA 2015, Lal 2008). The defining feature of these wetlands is the flooded soils. This is relevant to research since one of the primary indicators of microbial growth is the moisture of the soil; wetter soils are correlated with higher microbial biomass (Fierer. 2017). In particular, wetlands are efficient in storing carbon because the decomposition by microbes happens slower in this environment due to flooded conditions supporting anaerobic respiration which occurs at a slower rate compared to aerobic respiration. The dynamic hydrology of wetland ecosystems makes them important control points for microbial-climate change feedback (Oddershede 2018).

Bacterial diversity and community composition influence plant growth, showing an importance for more research in examining environmental changes affecting microbes in response to nutrient additions (Bastida et al. 2021). This study addresses the question, how does long term fertilization interact with hydrologic status to influence wetland plant and microbial biomass and bacterial community composition? First, we hypothesized that if fertilization increases plant biomass which, in turn, provides more carbon resources into soils, then microbial biomass will increase since there will be more available energy to support increased growth rates. Second, we hypothesized that if the plant rhizosphere provides organic carbon and oxygen to the soil environment, then microbial biomass will relate to belowground plant biomass more than aboveground plant biomass. Third, we hypothesized that if nutrient enrichment adds previously unavailable nutrients to the soil, then microbial diversity will also increase since fertilization provides more nutrients to support bacterial growth. To test these hypotheses, we collected soil cores and plant litter samples from a long-term wetland ecology fertilization experiment. We focused on areas that were mowed to simulate historical fire-maintained plant communities. At this site, we measured soil carbon and plant biomass and characterized the bacterial community using 16S rRNA targeted amplicon sequencing. We also identified bacterial taxa from the soil cores that are putatively considered oligotrophic or copiotrophic based on previous studies. This ongoing work provides insight into how nutrient enrichment modifies wetland carbon storage potential.

## METHODS

### Experimental design

For this study, we used data collected from a long-term ecological experiment on East Carolina University’s West research campus. This long-term ecological experiment studies factors that affect both the plant and microbial communities of wetlands. In particular, the experiment focuses on the effects of mowing fertilization in wetland communities. Detailed descriptions of the characteristics of the study site and experimental design can be found in Goodwillie et al. (2020). Briefly, this experiment represents a 2 × 2 factorial design where two levels of mowing (mowed, unmowed) are crossed with two levels of fertilization (fertilized, unfertilized) and replicated across eight blocks (Figure 1). All plots were 20 × 30 m blocks and treatments included nitrogen-phosphorus-potassium (N-P-K) 10-10-10 pellet fertilizer applied three times per year in February, June, and October (Figure 1). For plant diversity and soil sampling, three designated locations within each plot were sampled annually. Half of the plots are located adjacent to a ditch (drier soils) compared to away from the ditch, where soil conditions are wetter (Goodwillie et al. 2020). We focused on data collected from the unfertilized/mowed and fertilized/mowed plots because these plots are more representative of how this wetland was naturally maintained by wildfires before human intervention (Goodwillie et al. 2020).

**Figure 1.**
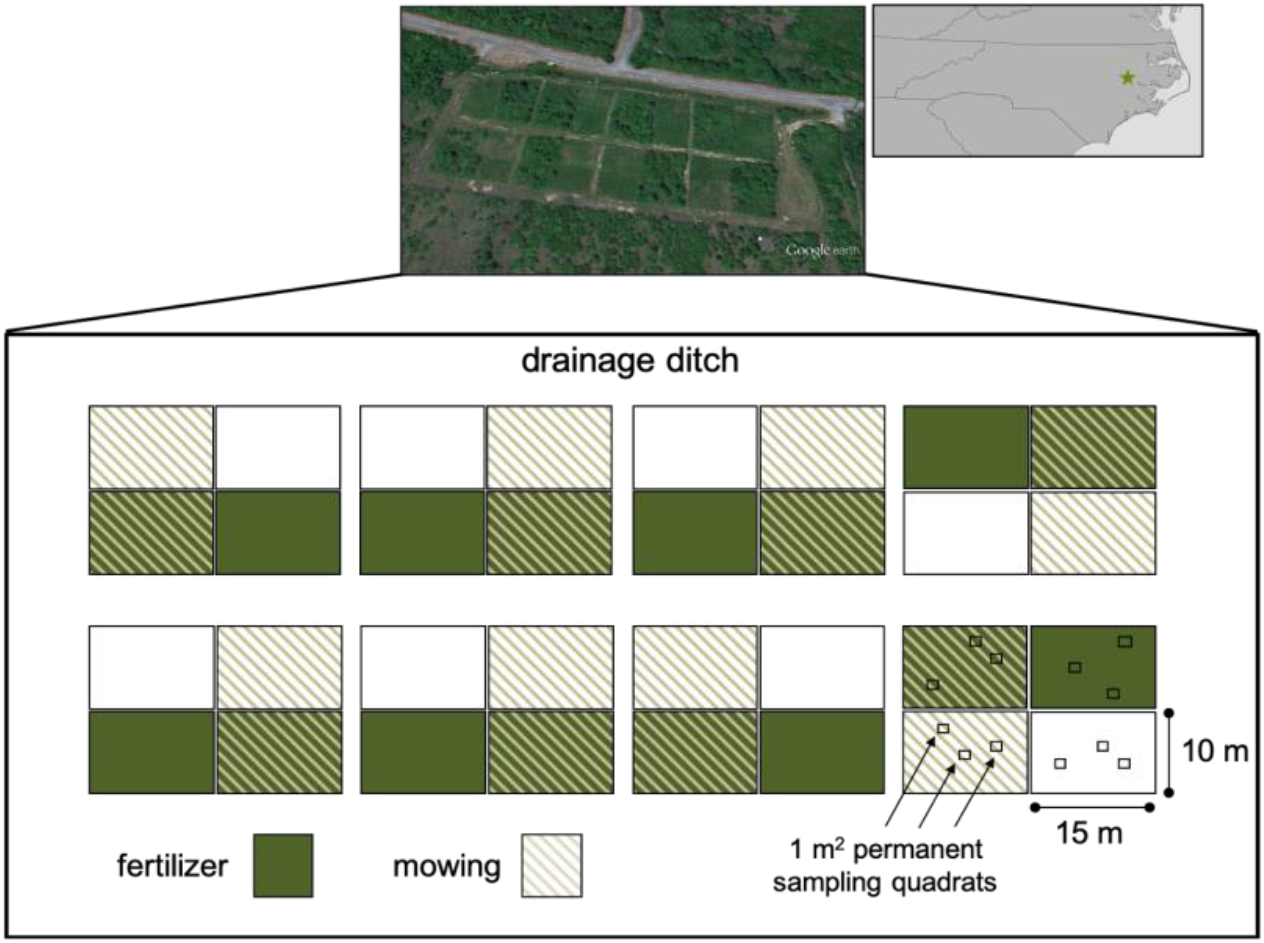
Experimental design of a long-term ecological experiment to test the effects of fertilizer and disturbance by mowing on plant and microbial communities at East Carolina University’s West Research Campus (WRC), Greenville, North Carolina, USA. Fertilizer application 3x a year with 10-10-10 NPK. Mowing and raking 1x a year.

### Plant and microbial biomass

To address the impacts of fertilization on plant and microbial biomass, we analyzed soil cores and plant litter. We collected and clipped plant litter samples in a 0.25-m^2^ area at ground level from mowed and mowed/fertilized plots after the growing season in 2010 and 2017. We then dried the samples at 60 °C for 48 hours and weighed the above ground plant biomass. We collected the biomass data at three randomly chosen locations per plot for a total of 48 samples according to previously described methods (Goodwillie et al. 2020). We collected the soil core for belowground root biomass using a PVC tube (8 cm diameter, 15 cm depth). Then, we dry sieved the soil sample to pick out the plant roots. We dried the roots in an oven at 60 °C for 48 hours and then weighed them. To calculate microbial biomass, we measured carbon using a chloroform fumigation method (McDaniel and Grandy 2016). We measured 5g of each soil sample into two 50 mL tubes. We treated one 5g sample with chloroform and the other was a control with no additions. The chloroform incubation occurred overnight to kill living cells. We collected extractable carbon and nitrogen from the soil and then measured for dissolved carbon and nitrogen. We calculated the microbial biomass from the difference in dissolved carbon and nitrogen between fumigated and non-fumigated soils.

### Bacterial community composition

We extracted DNA from each soil sample using the Qiagen DNeasy PowerSoil kit. After extraction, we used these sample DNA as a template in PCRs with a bacterial 515F/806RB barcoded primer set originally developed by the Earth Microbiome Project to target the V4 region of the bacterial 16S subunit of the rRNA gene (Caporaso et al. 2012, Koceja et al. 2021, Bledsoe et al. 2020). For each of the samples, we prepared three 50 μL PCR libraries by combining 35.75 μL molecular grade water, 5 μL Amplitaq Gold 360 10× buffer, 5 μL MgCl2 (25 mmol/L), 1 μL dNTPs (40 mmol/L total, 10 mmol/L each), 0.25 μL Amplitaq Gold poly-merase, 1 μL 515 forward barcoded primer (10 μmol/L), 1 μL 806 reverse primer (10 μmol/L), and 1 μL DNA template (10 ng/μL). The thermocycler conditions for PCRs were as follows: initial denaturation (94°C, 3 min): 30 cycles of 94°C for 45s, 50°C for 30s, and 72°C for 90s; final elongation (72°C, 10 min). We combined the Triplicate PCR reactions for each sample and cleaned them according to the Axygen AxyPrep Magnetic Bead Purification Kits protocol (Corning Life Sciences). Following cleaning, we quantified PCR products using Quant-iT dsDNA BR (broad-range) assay (Thermo Scientific, Waltham, Massachusetts, USA). We pooled libraries in an equimolar concentration of 5 ng/μL after being diluted to a concentration of 10 ng/ μL. The Indiana University Center for Genomics and Bioinformatics sequenced the pooled libraries using the Illumina MiSeq platform using paired-end reads (Illumina Reagent Kit v2, 500 reaction kit). Following sequencing, we processed raw sequences using a standard mothur pipeline (v1.40.1; Schloss et al. 2009, Kozich et al. 2013). We assembled contigs from paired end reads and quality trimmed using a moving average quality score (minimum score of 35 bp). Sequences were aligned to the SILVA rRNA gene database (v.128; Quast et al. 2013), and chimeric sequences were removed using the VSEARCH algorithm (Rognes et al. 2016). Formation of operational taxonomic units (OTUs) involved dividing sequences based on taxonomic class and then binned into OTUs with a 97% sequence similarity level. Operational taxonomic units were classified using the SILVA rRNA database (v128; Yilmaz et al. 2014, Glöckner et al. 2017). We used *vegan::diversity* to calculate bacterial species diversity by calculating Shannon diversity index 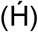. These methods are also described in Koceja et al. 2021 and Bledsoe et al. 2020.

### Statistical analyses

All statistical calculations were done in the R environment (R v4.1.3, R Core Team 2022) and graphs were constructed using the ggplot2 package. We visualized bacterial community composition using principal-coordinate analysis (PCoA) based on Bray-Curtis dissimilarity matrix to illustrate bacterial community responses to nutrient enrichment. We tested for differences in bacterial community composition among treatments using permutational multivariate analysis of variance (PERMANOVA). We used the *vegan::adonis* function to perform hypothesis testing using PERMANOVA (Bledsoe et al. 2020). We subsetted common bacterial taxa identified to be either oligotrophic or copiotrophic to test the effects of fertilization on life-history strategy. We visualized patterns in copiotroph and oligotroph communities using a PCoA. Lastly, we identified which bacterial taxa to individually graph that would most accurately represent changes of the copiotroph or oligotroph communities with Dufrene-Legendre indicator species analysis using the *labdsv::indval* function.

## RESULTS

### Soil carbon and plant productivity

Ditch and fertilization effects influenced soil environmental factors and plant biomass to different degrees. Ditch and fertilization effects greatly influenced wetland conditions, where wet conditions tended to have higher carbon concentrations compared to drier, ditch conditions. However, soil carbon concentrations were similar in fertilized and unfertilized treatment plots (ANOVA, treatment, F_1, 6_ = 0.862, P = 0.389, Figure 2). Results show an overall non-significant increase in soil carbon (ANOVA, treatment:ditch, F_1, 6_ = 0.009, P = 0.929, Figure 2). Aboveground plant biomass showed a significant increase in response to the addition of fertilizer, especially in the wet ditch plots (ANOVA, treatment:ditch, F_1, 38_ = 11.02, P = 0.002, Figure 3A). In contrast, fertilization did not influence belowground plant biomass (ANOVA, treatment, F_1, 38_ = 1.97, P = 0.168, Figure 3B).

**Figure 2:**
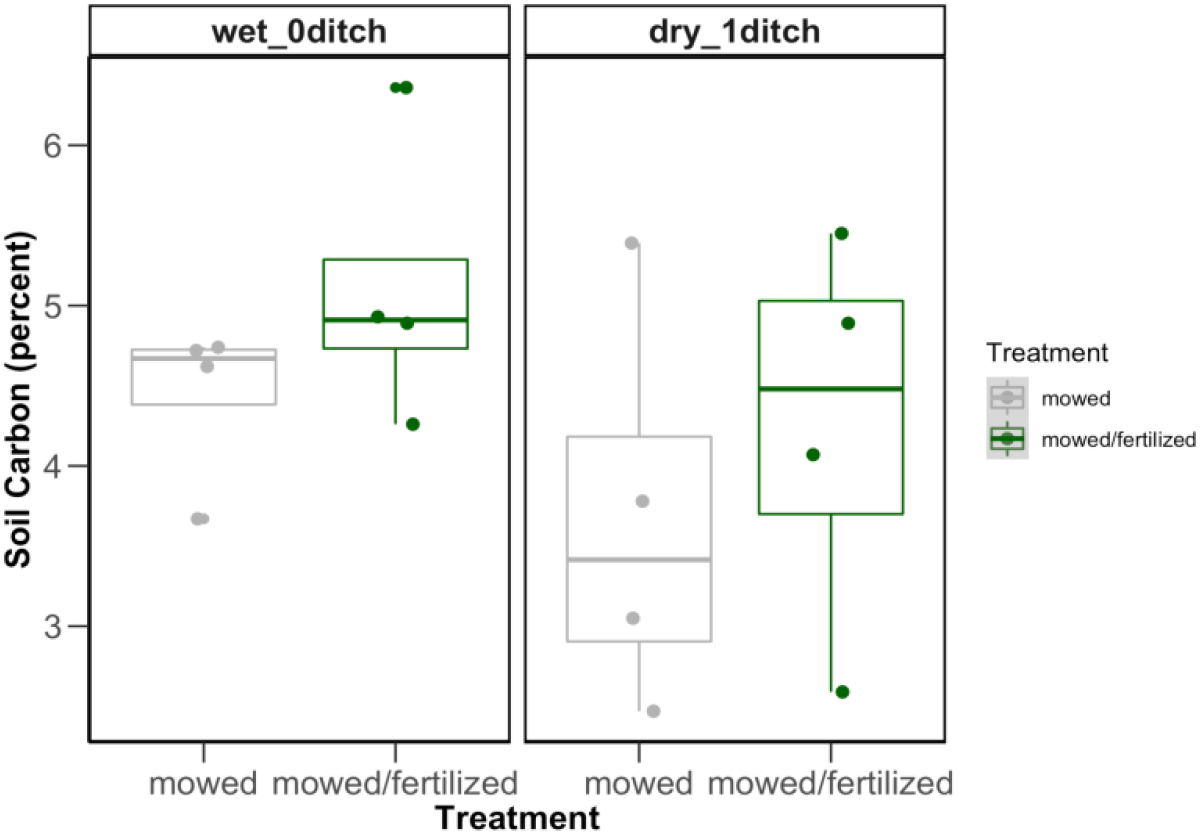
Boxplots representing soil carbon measured as a percent of total soil composition in fertilized and unfertilized plots. Boxplots and symbols are colored according to fertilization treatment (gray = unfertilized, green = fertilized) at wetter mowed plots situated away from the drainage ditch (left) compared to drier mowed plots situated close to the drainage ditch (right). The boxplot is a visual representation of 5 key summary statistics: the median, the 25% and 75% percentiles, and the whiskers which represent the feasible range of the data as determined by +/- 1.5 × the interquartile range. Symbols represent data points for individual plot samples.

**Figure 3:**
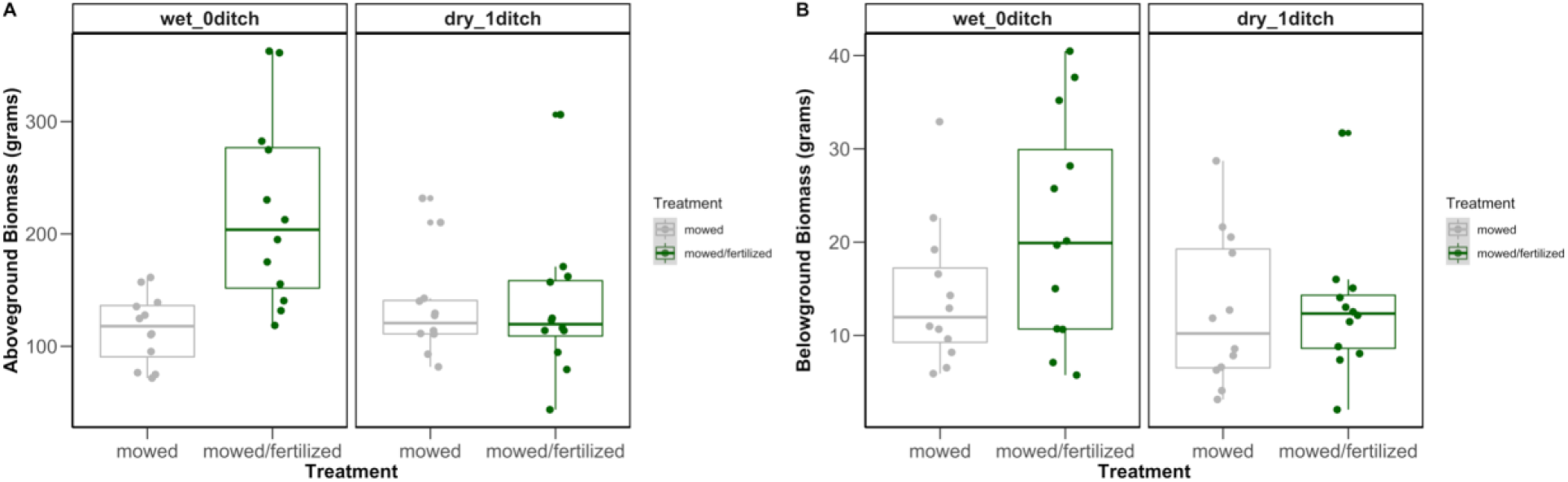
Boxplots representing aboveground (A) and belowground (B) biomass changes due to fertilization and ditch effects. Boxplots and symbols are colored according to fertilization treatment (gray = unfertilized, green = fertilized) at wetter mowed plots situated away from the drainage ditch (left panel) compared to drier mowed plots situated close to the drainage ditch (right panel). The boxplot is a visual representation of 5 key summary statistics: the median, the 25% and 75% percentiles, and the whiskers which represent the feasible range of the data as determined by +/- 1.5 × the interquartile range. Symbols represent data points for individual plot samples.

### Microbial biomass and diversity

Fertilization was shown to have varying degrees of impacts on variables related to microbial life. Microbial biomass, which was measured as total organic carbon of each soil sample, was similar in each plot and was not significantly changed by ditch effects alone (ANOVA, ditch, F_1, 6_ = 0.006, P = 0.941, Figure 4). After fertilization, microbial biomass decreased slightly in wet ‘no ditch’ conditions but increased in dry ditch conditions (ANOVA, treatment, F_3, 66_ = 4.90, P = 0.004, Figure 4A). In addition, bacterial Shannon diversity greatly increased due to fertilization (ANOVA, treatment, F_1, 6_ = 18.9, P = 0.005 Figure 4B).

**Figure 4:**
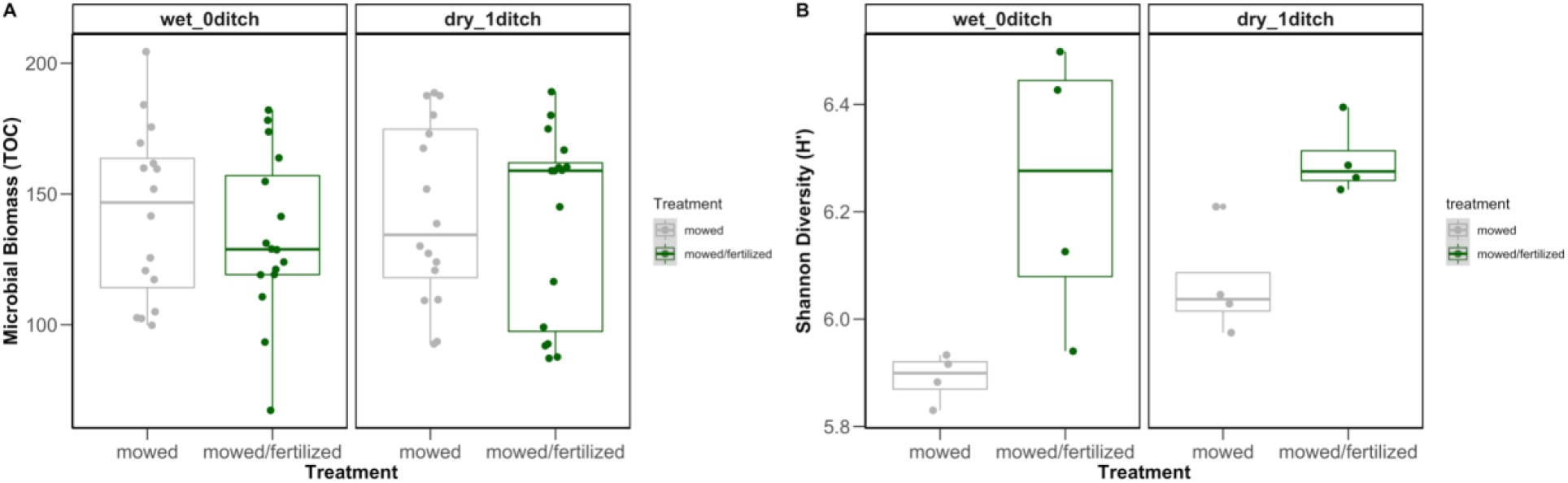
Boxplots representing microbial biomass (A) and bacterial Shannon Diversity Index (H’) in fertilized and unfertilized plots. Boxplots and symbols are colored according to fertilization treatment (gray = unfertilized, green = fertilized) at wetter mowed plots situated away from the drainage ditch (left) compared to drier mowed plots situated close to the drainage ditch (right). The boxplot is a visual representation of 5 key summary statistics: the median, the 25% and 75% percentiles, and the whiskers which represent the feasible range of the data as determined by +/- 1.5 × the interquartile range. Symbols represent data points for individual plot samples.

### Relationship between plant and microbial biomass

We determined the extent that plant biomass and microbial biomass are related. We examined ratios of microbial biomass to aboveground plant biomass (Figure 5A) and microbial biomass to belowground plant biomass (Figure 5B). There is no significant relationship between microbial biomass and aboveground plant biomass according to treatment (Pearson’s correlation, unfertilized: r = 0.040, P = 0.851, fertilized: r = -0.135, P = 0.549; Fig. 5A). Results also showed there was no significant correlation between belowground plant biomass to microbial biomass (Pearson’s correlation, unfertilized: r = 0.343, P=0.101, fertilized: r=-0.175, P=0.413; Fig. 5B). But an interesting positive trend was observed for microbial biomass and belowground biomass in the unfertilized plots (Figure 5B).

**Figure 5:**
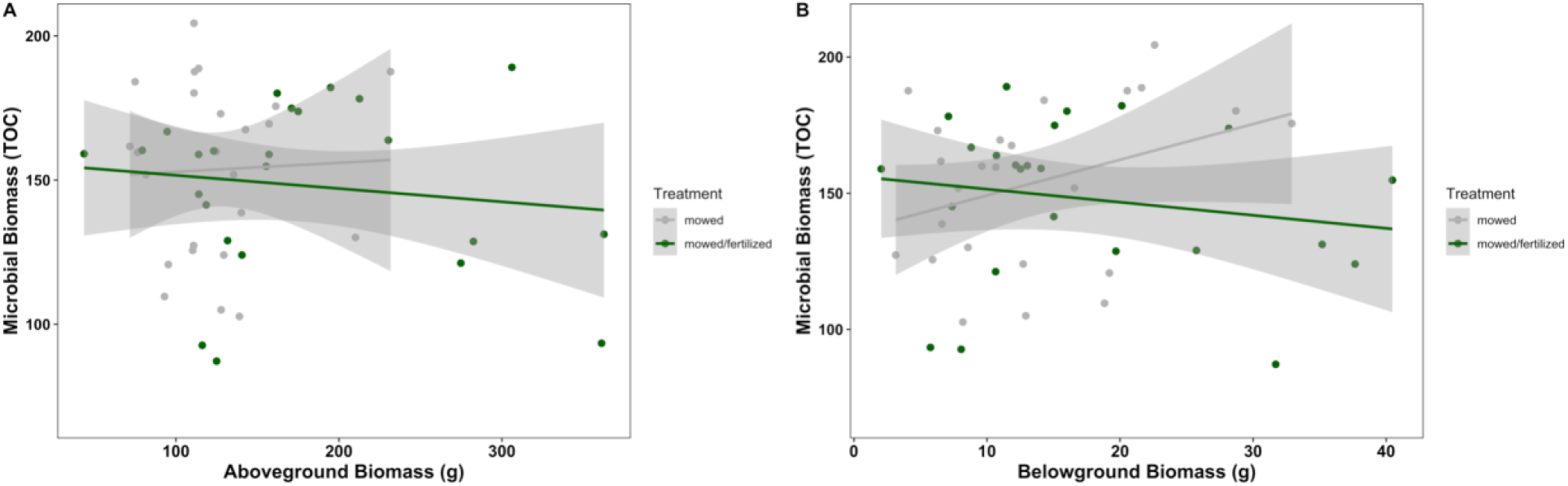
Scatter plot showing correlation between belowground plant biomass and microbial biomass along with aboveground plant biomass and microbial biomass. Plots and symbols are colored according to fertilization treatment (gray = unfertilized, green = fertilized) at wetter mowed plots situated away from the drainage ditch (left) compared to drier mowed plots situated close to the drainage ditch (right). Symbols represent data points for individual plot samples.

### Microbial communities and OTU-level patterns

Results showed significant differences in microbial community composition due to fertilization effects. Specifically, bacterial taxa were categorized into oligotrophs and copiotrophs to examine a subset of the bacterial community. Ordination plots for oligotrophic taxa showed significant changes in community composition of Planctomycetes (PERMANOVA, treatment, R^2^ = 0.268, P = 0.001, Figure 6A) and Verrucomicrobia (PERMANOVA, treatment, R^2^ = 0.217, P = 0.006, Figure 6B). The determined species indicator for Planctomycetes had a notable decrease in relative abundance after fertilization (ANOVA, treatment, F_1, 6_ =16.3, P = 0.007, Figure 7A). Similar results were observed for the Verrucomicrobia species indicator where relative abundance was higher in fertilized compared to unfertilized plots (ANOVA, treatment, F = 15.851, P = 0.007, Figure 7B). Ordination plots for copiotrophic taxa revealed distinct bacterial communities according to treatment for Rhizobiales and ditch effect for Bacillales. Bacillales community composition showed less change after treatment (PERMANOVA, treatment, R^2^ = 0.073, P = 0.273, Figure 8A) compared to the significant change in Rhizobiales community composition (PERMANOVA, treatment, R^2^ = 0.271, P = 0.001, Figure 8B). Further, Bacillales community composition was influenced by ditch effects (PERMANOVA, ditch, R^2^=0.125, P=0.074). However, species indicator taxon showed no significant change in relative abundance for Bacillales (ANOVA, treatment, F_1, 6_ =0.234, P= 0.646, Figure 9A) or Rhizobiales (ANOVA, treatment, F_1, 6_ = 3.13, P = 0.127, Figure 9B) between fertilized and unfertilized plots.

**Figure 6:**
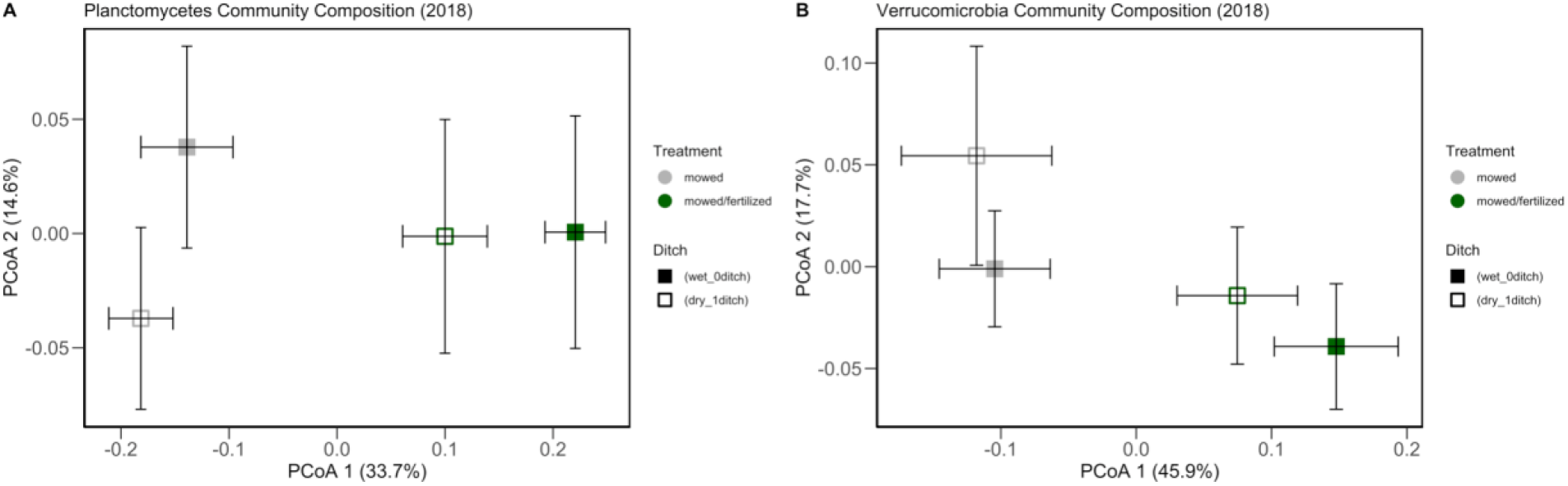
Ordination plots representing community composition of **oligotrophic** taxa in fertilized and unfertilized plots during the year 2018. These ordination plots display average composition from four replicate field samples. Gray boxes show unfertilized and green show fertilized. Groups that are more similar in community composition are closer together than groups with points that are far apart.

**Figure 7:**
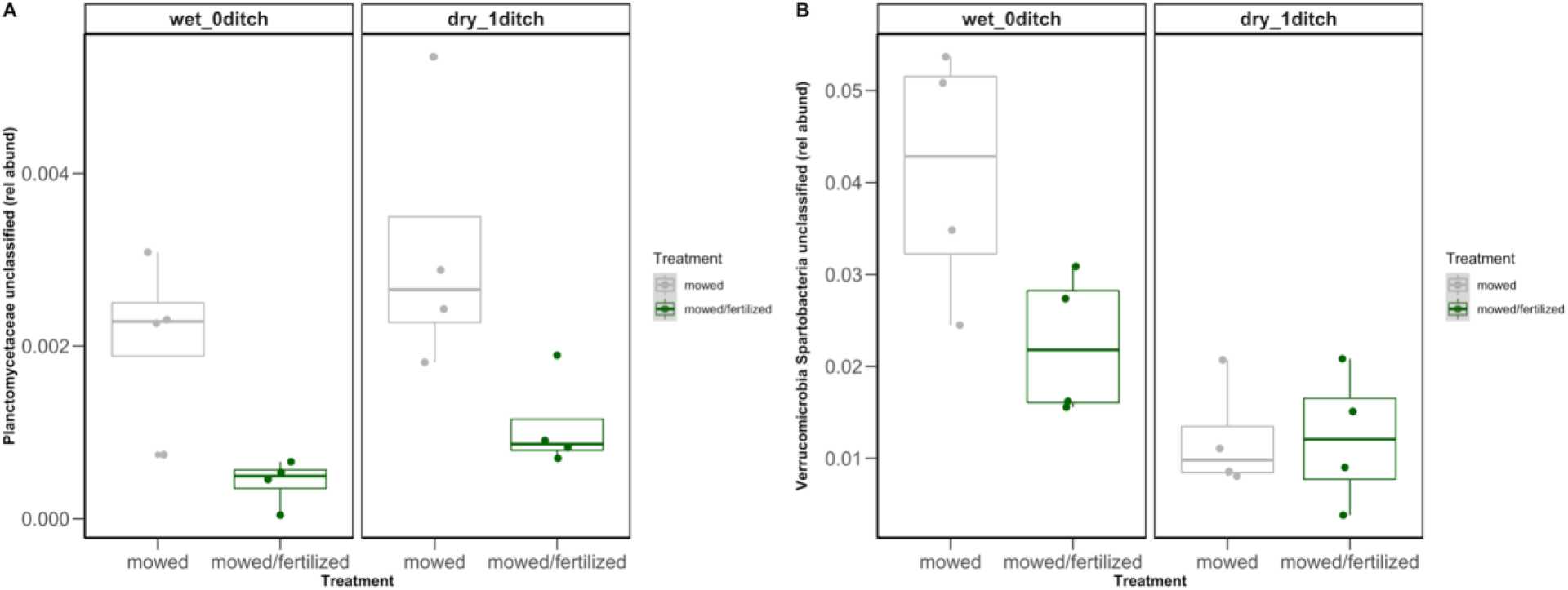
Boxplots representing relative abundance of **oligotrophic** taxa in fertilized and unfertilized plots. Boxplots and symbols are colored according to fertilization treatment (gray = unfertilized, green = fertilized) at wetter mowed plots situated away from the drainage ditch (left) compared to drier mowed plots situated close to the drainage ditch (right). The boxplot is a visual representation of 5 key summary statistics: the median, the 25% and 75% percentiles, and the whiskers which represent the feasible range of the data as determined by +/- 1.5 × the interquartile range. Symbols represent data points for individual plot samples.

**Figure 8:**
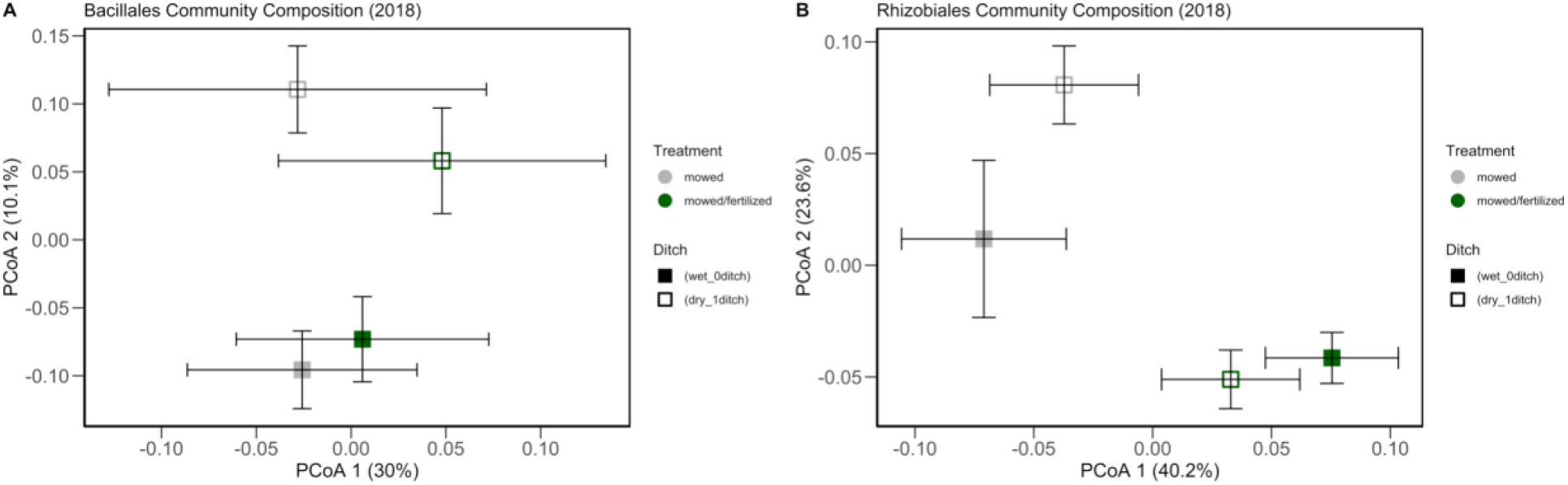
Ordination plots representing community composition of **copiotrophic** taxa in fertilized and unfertilized plots during the year 2018. These ordination plots display average composition from four replicate field samples. Gray boxes show unfertilized and green show fertilized. Groups that are more similar in community composition are closer together than groups with points that are far apart.

**Figure 9:**
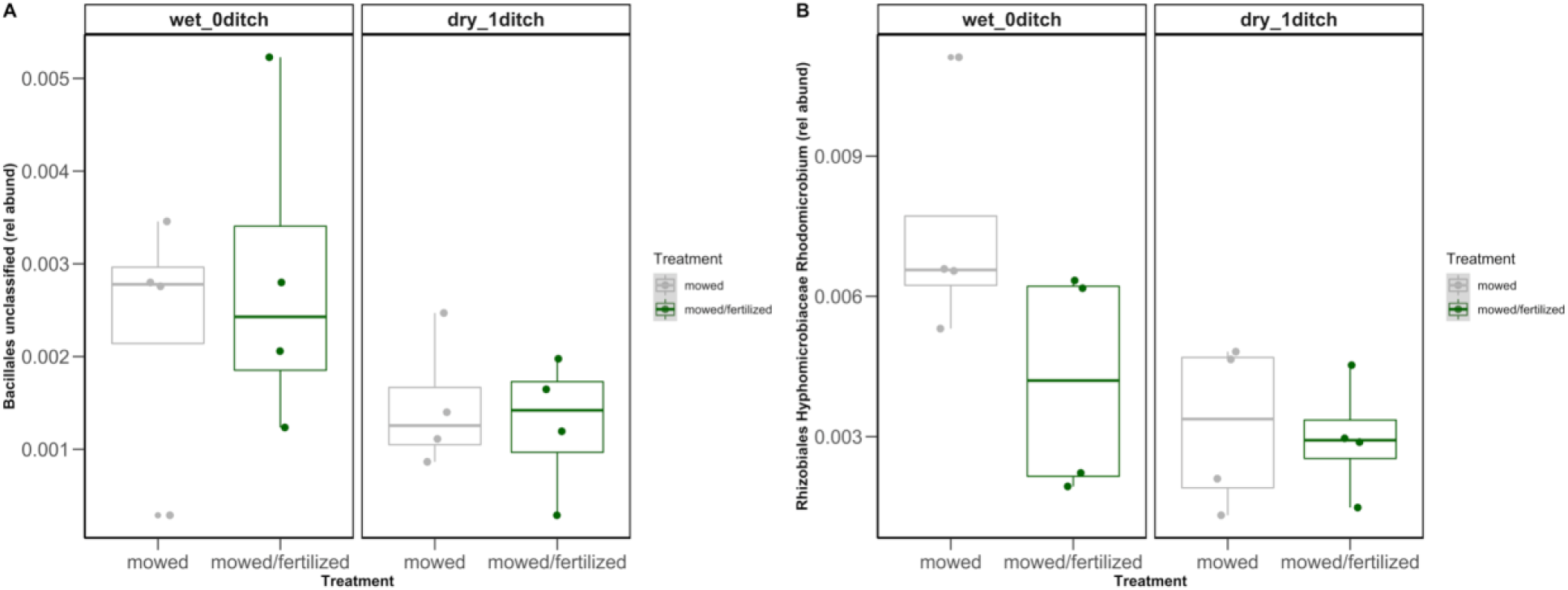
Boxplots representing relative abundance of **copiotrophic** taxa in fertilized and unfertilized plots. Boxplots and symbols are colored according to fertilization treatment (gray = unfertilized, green = fertilized) at wetter mowed plots situated away from the drainage ditch (left) compared to drier mowed plots situated close to the drainage ditch (right). The boxplot is a visual representation of 5 key summary statistics: the median, the 25% and 75% percentiles, and the whiskers which represent the feasible range of the data as determined by +/- 1.5 × the interquartile range. Symbols represent data points for individual plot samples.

## DISCUSSION

Long-term nutrient additions to low nutrient coastal plain wetland communities are showing signs of accelerated soil carbon losses with fertilization. Fertilization effects are differentially influencing plant biomass, microbial biomass, and bacterial community composition. Despite continued increases in plant aboveground biomass; belowground plant biomass, microbial biomass carbon, and total organic carbon were similar in fertilized plots compared to unfertilized while soil moisture continues to play an interesting role in accelerating carbon content loss. Soil carbon concentration was similar in fertilized plots compared to unfertilized plots. In contrast, previous studies found an initial increase in soil carbon due to fertilization (Jesmin et al. 2021, Maltas et al. 2018). While nutrient addition did not influence soil carbon concentration, wetter soil conditions were associated with higher soil carbon content. Though wet soil conditions support slowed decomposition rates, fertilization effects can compound soil carbon losses. Carbon storage is expected to decrease in coming years due to the short turnover of the newly added carbon from fertilization (Tipping et al. 2017). The increase of nutrients associated with increased carbon in the soil also enhances plant growth (Goodwillie et al. 2020).

In this experiment, aboveground plant biomass increased due to fertilization. However, belowground plant biomass did not change due to the fertilization treatment. Fertilization influences microbial biomass in differing ways depending on hydrologic status (i.e., wetter soil conditions away from a drainage ditch versus drier soil conditions by a drainage ditch). In the wet plots, microbial biomass increased slightly while in dry plots there was a decrease in biomass. This coincides with the lack of change in belowground biomass as the rhizosphere is where we sampled to determine microbial biomass. With the overall surface area of the roots unchanged (i.e. similar belowground biomass measured), soil moisture compared to plant root inputs likely plays a stronger role in controlling fertilization effects on microbial processes (Fierer 2017). Together, these results support our first hypothesis that increasing plant biomass will also increase microbial biomass.

There have been extensive studies that detail the effects of fertilization and nutrient addition on plant biomass and microbial communities as we have investigated in this experiment (Harpole et al. 2016, Bledsoe et al. 2020) along with studies to explain the relationships that cause these plants and microbes to be potentially correlated. We measured the relationship between plant biomass and microbial biomass. Results revealed no relationship between plant biomass (above or belowground) and soil microbial biomass. These findings suggest that the overall biomass of plant life does not directly influence the amount of microbial biomass found in samples. These results do not support the second hypothesis of belowground plant biomass having more of a correlation to microbial biomass than aboveground plant biomass is inconclusive, but there is an emerging positive relationship between microbial biomass and belowground plant biomass for unfertilized plots only. Our results could have been limited by an insufficient sample size along with disregarding the ditch effects. Increasing the amount of data we examine may provide different results. In a previous study, microbial biomass was correlated with belowground plant biomass (Gutiérrez-Girón et al. 2014). Future studies would also benefit from evaluating nutrient enrichment effects on bacterial vs. fungal biomass and the association with plant biomass.

The final focus of this study was to determine the effects of fertilization on bacterial diversity and community composition. We found that bacterial Shannon diversity significantly increased due to fertilization, similar to previous studies (Bledsoe et al. 2020, Li et al. 2021, Koceja et al. 2021). In contrast, other studies found either no change or a decrease in bacterial diversity following fertilization due to the coinciding decrease in plant diversity or soil pH (Ren et al. 2020, Feng et al. 2021). The increase in diversity found in this study is likely motivated by the decrease in competition over nutritional resources in fertilized plots. Soil moisture also has a large effect on microbial diversity since wetter, more irrigated areas show a larger increase in diversity than in dry areas (Li et al. 2021). This increase of bacterial diversity is also shown to increase the rate of decomposition in the soil (Lange et al 2015, Koceja et al. 2021).

Soil bacterial life history strategy can directly influence decomposition rates. Slow-growing microbes are more efficient at using soil carbon and decompose matter slower than their fast-growing counterparts (Roller and Schmidt 2015). As a consequence, slower decomposition will release less carbon into the atmosphere and would positively contribute to the soil’s ability to store carbon (Fierer et al. 2007). This study focused on oligotrophic and copiotrophic bacteria to explore nutrient effects on soil bacterial life history. Previous studies categorize the phyla of Planctomycetes and Verrucomicrobia to be oligotrophic (Ho et al. 2017, Pepe-Ranney et al. 2016). The orders of Bacialles and Rhizobiales were categorized as copiotrophic (Ho et al. 2017, Cleveland et al. 2007). By focusing on OTU’s classified as Planctomycetes or Verrucomicrobia, we examined fertilization and soil moisture effects on specific subsets of the bacterial community. Further, results showed that oligotrophic bacteria relative abundance decreased due to fertilization while copiotrophic bacteria relative abundances were not significantly affected. The relative abundances of oligotrophic bacterial taxa were more negatively affected by fertilization than the copiotrophic bacterial taxa. In a non-nutrient limiting environment, copiotrophs can use the available nutrients more quickly than the oligotrophs and outcompete them in the process (Roller and Schmidt 2015, Fierer et al. 2007). The decrease in oligotroph relative abundance may have negative impacts on wetland carbon storage since they are being replaced by bacteria and other microorganisms that likely respire carbon at higher rates. The extent of influence that bacterial life history strategy has on carbon storage potential is unknown but oligotrophic bacteria are known to have less metabolic activity compared to copiotrophic bacteria leading to increased carbon use efficiency in oligotrophs (Fierer et al. 2007, Ho et al. 2017).

Despite the decrease in oligotroph relative abundance, the fertilization did not affect the relative abundance of the copiotrophic taxa. There are many possible reasons for the mixed results relating to the copiotrophic bacteria composition and relative abundance. One possibility is that although the bacteria are classified as copiotrophs, their resource requirements differ from the average bacteria in that category. Rhizobiales, for instance, are nitrogen-fixing bacteria that have special characteristics to allow for enhanced symbiosis with plant roots that many other copiotrophic bacteria do not have (Garrido-Oter et al. 2018). Another reason could be that the taxa that were examined have characteristics of both oligotrophs and copiotrophs. Some copiotrophic bacteria, such as Actinobacteria, are able to break down complex carbon substrates that oligotrophic bacteria are more likely to process (Barder and Crawford 1981, Ho et al. 2017). Determining the life history strategy of soil bacteria is an important indicator of microbial-climate change effects. Future research should consider mechanisms according to life history traits.

## CONCLUSION

In this study, nutrient enrichment influenced plant growth and bacterial community composition of wetland habitats. Aboveground plant biomass, microbial biomass, and microbial diversity increased in response to fertilization while soil carbon and below ground biomass showed no changes. These results indicate that wetlands are experiencing ongoing nutrient-mediated changes in complex, interacting ways. In particular, long-term nutrient additions are differentially affecting bacterial community composition in ways that could diminish the slow growing taxa. The decreasing trend in oligotrophic bacteria abundance is expected to diminish wetland carbon storage potential as bacterial communities shift to respire carbon stored in the soil more quickly. This study reveals unexpected consequences of nutrient feedbacks on carbon storage due to interactive effects of soil moisture and fertilization on plant biomass and soil bacterial communities. In order to better protect the health of soil, we should further investigate the changes happening in wetland communities, especially the microbes within them as they are major contributors to global carbon storage. By characterizing the shifts happening to bacteria within wetland soil, we can counteract the effects of atmospheric deposition and direct additions of nutrients into the environment.

## ACKNOWLEDGEMENTS

We thank Allison Fisk, Emma Richards, Tom Vogel, Suelen Tullio, and Adam Gold for laboratory and field assistance in processing the 2018 samples. We thank Daniya Stephens, Aied Garcia, Tom Vogel, Joseph Whittington for laboratory and field assistance in processing the 2020 samples. We thank Carol Goodwillie, John Stiller, John Gill, and the East Carolina University grounds crew for their tireless efforts in maintaining the long-term ecological experiment. This work was supported by the National Summer Undergraduate Research Project (NSURP) and the National Science Foundation (DEB 1845845 to ALP).

## REFERENCES

Barder, M. J., and D. L. Crawford. 1981. Effects of carbon and nitrogen supplementation on lignin and cellulose decomposition by a Streptomyces. Canadian Journal of Microbiology 27:859–863.

Bastida, F., D. J. Eldridge, C. García, G. Kenny Png, R. D. Bardgett, and M. Delgado-Baquerizo. 2021. Soil microbial diversity–biomass relationships are driven by soil carbon content across global biomes. The ISME Journal 15:2081–2091.

Blaser, M. J., Z. G. Cardon, M. K. Cho, J. L. Dangl, T. J. Donohue, J. L. Green, R. Knight, M. E. Maxon, T. R. Northen, K. S. Pollard, and E. L. Brodie. 2016. Toward a Predictive Understanding of Earth’s Microbiomes to Address 21st Century Challenges. mBio 7.

Bledsoe, R., C. Goodwillie, and A. Peralta. 2020. Long-Term Nutrient Enrichment of an Oligotroph-Dominated Wetland Increases Bacterial Diversity in Bulk Soils and Plant Rhizospheres. mSphere 5.

Borer, E. T., J. B. Grace, W. S. Harpole, A. S. MacDougall, and E. W. Seabloom. 2017. A decade of insights into grassland ecosystem responses to global environmental change. Nature Ecology & Evolution 1:1–7.

Caporaso, J. G., C. L. Lauber, W. A. Walters, D. Berg-Lyons, J. Huntley, N. Fierer, S. M. Owens, J. Betley, L. Fraser, M. Bauer, N. Gormley, J. A. Gilbert, G. Smith, and R. Knight. 2012. Ultra-high-throughput microbial community analysis on the Illumina HiSeq and MiSeq platforms. The ISME Journal 6:1621–1624.

Cavicchioli, R., W. J. Ripple, K. N. Timmis, F. Azam, L. R. Bakken, M. Baylis, M. J. Behrenfeld, A. Boetius, P. W. Boyd, A. T. Classen, T. W. Crowther, R. Danovaro, C. M. Foreman, J. Huisman, D. A. Hutchins, J. K. Jansson, D. M. Karl, B. Koskella, D. B. Mark Welch, J. B. H. Martiny, M. A. Moran, V. J. Orphan, D. S. Reay, J. V. Remais, V. I. Rich, B. K. Singh, L. Y. Stein, F. J. Stewart, M. B. Sullivan, M. J. H. van Oppen, S. C. Weaver, E. A. Webb, and N. S. Webster. 2019. Scientists’ warning to humanity: microorganisms and climate change. Nature Reviews Microbiology 17:569–586.

Cleveland, C. C., D. R. Nemergut, S. K. Schmidt, and A. R. Townsend. 2007. Increases in soil respiration following labile carbon additions linked to rapid shifts in soil microbial community composition. Biogeochemistry 82:229–240.

Davidson, E., M. David, J. Galloway, R. Haeuber, J. Harrison, R. Howarth, R. Lowrance, T. Nolan, J. Peel, R. Pinder, C. Snyder, A. Townsend, and M. Ward. 2012. Excess Nitrogen in the U.S. Environment: Trends, Risks, and Solutions. Issues in Ecology.

Feng, Y., M. Delgado-Baquerizo, Y. Zhu, X. Han, X. Han, X. Xin, W. Li, Z. Guo, T. Dang, C. Li, B. Zhu, Z. Cai, D. Li, and J. Zhang. 2021. Responses of Soil Bacterial Diversity to Fertilization Are Driven by Local Environmental Context Across China. Engineering.

Fierer, N. 2017. Embracing the unknown: disentangling the complexities of the soil microbiome. Nature Reviews Microbiology 15:579–590.

Fornara, D. A., and D. Tilman. 2012. Soil carbon sequestration in prairie grasslands increased by chronic nitrogen addition. Ecology 93:2030–2036.

Garrido-Oter, R., R. T. Nakano, N. Dombrowski, K. W. Ma, A. C. McHardy, and P. Schulze-Lefert. 2018. Modular Traits of the Rhizobiales Root Microbiota and Their Evolutionary Relationship with Symbiotic Rhizobia. Cell Host and Microbe 24:undefined-undefined.

Glöckner, F. O., P. Yilmaz, C. Quast, J. Gerken, A. Beccati, A. Ciuprina, G. Bruns, P. Yarza, J. Peplies, R. Westram, and W. Ludwig. 2017. 25 years of serving the community with ribosomal RNA gene reference databases and tools. Journal of Biotechnology 261:169– 176.

Goodwillie, C., and W. R. Franch. 2006. An experimental study of the effects of nutrient addition and mowing on a ditched wetland plant community: results of the first year. Journal of the North Carolina Academy of Science 122:106–117.

Goodwillie, C., M. W. McCoy, and A. L. Peralta. 2020. Long-term nutrient enrichment, mowing, and ditch drainage interact in the dynamics of a wetland plant community. Ecosphere 11:e03252.

Gutiérrez-Girón, A., A. Rubio, and R. G. Gavilán. 2014. Temporal variation in microbial and plant biomass during summer in a Mediterranean high-mountain dry grassland. Plant and Soil 374:803–813.

Harpole, W. S., L. L. Sullivan, E. M. Lind, J. Firn, P. B. Adler, E. T. Borer, J. Chase, P. A. Fay, Y. Hautier, H. Hillebrand, A. S. MacDougall, E. W. Seabloom, R. Williams, J. D. Bakker, M. W. Cadotte, E. J. Chaneton, C. Chu, E. E. Cleland, C. D’Antonio, K. F. Davies, D. S. Gruner, N. Hagenah, K. Kirkman, J. M. H. Knops, K. J. La Pierre, R. L. McCulley, J. L. Moore, J. W. Morgan, S. M. Prober, A. C. Risch, M. Schuetz, C. J. Stevens, and P. D. Wragg. 2016. Addition of multiple limiting resources reduces grassland diversity. Nature 537:93–96.

Ho, A., D. P. Di Lonardo, and P. L. E. Bodelier. 2017. Revisiting life strategy concepts in environmental microbial ecology. FEMS Microbiology Ecology 93.

Hoosbeek, M., M. Lukac, D. Dam, D. Godbold, E. Velthorst, F. A. Bondi, A. Peressotti, M. F. Cotrufo, P. Angelis, and G. Scarascia-Mugnozza. 2004. More new carbon in the mineral soil of a poplar plantation under Free Air Carbon Erichment (POPFACE): Cause of increased priming effect? Global Biogeochemical Cycles 18.

Jesmin, T., D. T. Mitchell, and R. L. Mulvaney. 2021. Short-Term Effect of Nitrogen Fertilization on Carbon Mineralization during Corn Residue Decomposition in Soil. Nitrogen 2:444–460.

Jiang, D., L. Chen, N. Xia, E. Norgbey, D. A. Koomson, and W. K. Darkwah. 2020. Elevated atmospheric CO2 impact on carbon and nitrogen transformations and microbial community in replicated wetland. Ecological Processes 9:57.

Knight, R., A. Vrbanac, B. C. Taylor, A. Aksenov, C. Callewaert, J. Debelius, A. Gonzalez, T. Kosciolek, L.-I. McCall, D. McDonald, A. V. Melnik, J. T. Morton, J. Navas, R. A. Quinn, J. G. Sanders, A. D. Swafford, L. R. Thompson, A. Tripathi, Z. Z. Xu, J. R. Zaneveld, Q. Zhu, J. G. Caporaso, and P. C. Dorrestein. 2018. Best practices for analysing microbiomes. Nature Reviews Microbiology 16:410–422.

Koceja, M. E., R. B. Bledsoe, C. Goodwillie, and A. L. Peralta. 2021. Distinct microbial communities alter litter decomposition rates in a fertilized coastal plain wetland. Ecosphere 12:e03619.

Kozich, J. J., S. L. Westcott, N. T. Baxter, S. K. Highlander, and P. D. Schloss. 2013. Development of a Dual-Index Sequencing Strategy and Curation Pipeline for Analyzing Amplicon Sequence Data on the MiSeq Illumina Sequencing Platform. Applied and Environmental Microbiology 79:5112–5120.

Kuzyakov, Y. 2010. Priming effects: Interactions between living and dead organic matter. Soil Biology and Biochemistry 42:1363–1371.

Lal, R. 2008. Carbon sequestration. Philosophical Transactions of the Royal Society B: Biological Sciences 363:815–830.

Lange, M., N. Eisenhauer, C. A. Sierra, H. Bessler, C. Engels, R. I. Griffiths, P. G. Mellado-Vázquez, A. A. Malik, J. Roy, S. Scheu, S. Steinbeiss, B. C. Thomson, S. E. Trumbore, and G. Gleixner. 2015. Plant diversity increases soil microbial activity and soil carbon storage. Nature Communications 6:6707.

Li, H., H. Wang, B. Jia, D. Li, Q. Fang, and R. Li. 2021. Irrigation has a higher impact on soil bacterial abundance, diversity and composition than nitrogen fertilization. Scientific Reports 11:16901.

Maltas, A., H. Kebli, H. R. Oberholzer, P. Weisskopf, and S. Sinaj. 2018. The effects of organic and mineral fertilizers on carbon sequestration, soil properties, and crop yields from a long-term field experiment under a Swiss conventional farming system. Land Degradation & Development 29:926–938.

McDaniel, M. D., and A. S. Grandy. 2016. Soil microbial biomass and function are altered by 12 years of crop rotation. SOIL 2:583–599.

Oddershede, A., C. Violle, A. Baattrup-Pedersen, J.-C. Svenning, and C. Damgaard. 2019. Early dynamics in plant community trait responses to a novel, more extreme hydrological gradient. Journal of Plant Ecology 12:327–335.

Pepe-Ranney, C., A. N. Campbell, C. N. Koechli, S. Berthrong, and D. H. Buckley. 2016. Unearthing the Ecology of Soil Microorganisms Using a High Resolution DNA-SIP Approach to Explore Cellulose and Xylose Metabolism in Soil. Frontiers in Microbiology 7.

Quast, C., E. Pruesse, P. Yilmaz, J. Gerken, T. Schweer, P. Yarza, J. Peplies, and F. O. Glöckner. 2013. The SILVA ribosomal RNA gene database project: improved data processing and web-based tools. Nucleic Acids Research 41:D590–D596.

Ramirez, K. S., J. M. Craine, and N. Fierer. 2012. Consistent effects of nitrogen amendments on soil microbial communities and processes across biomes. Global Change Biology 18:1918–1927.

Ren, N., Y. Wang, Y. Ye, Y. Zhao, Y. Huang, W. Fu, and X. Chu. 2020. Effects of Continuous Nitrogen Fertilizer Application on the Diversity and Composition of Rhizosphere Soil Bacteria. Frontiers in Microbiology 11.

Rognes, T., T. Flouri, B. Nichols, C. Quince, and F. Mahé. 2016. VSEARCH: a versatile open source tool for metagenomics. PeerJ 4:e2584.

Roller, B. R. K., and T. M. Schmidt. 2015. The physiology and ecological implications of efficient growth. The ISME journal 9:1481–1487.

Schloss, P. D., S. L. Westcott, T. Ryabin, J. R. Hall, M. Hartmann, E. B. Hollister, R. A. Lesniewski, B. B. Oakley, D. H. Parks, C. J. Robinson, J. W. Sahl, B. Stres, G. G. Thallinger, D. J. Van Horn, and C. F. Weber. 2009. Introducing mothur: open-source, platform-independent, community-supported software for describing and comparing microbial communities. Applied and Environmental Microbiology 75:7537–7541.

Schwede, D. B., and G. G. Lear. 2014. A novel hybrid approach for estimating total deposition in the United States. Atmospheric Environment 92:207–220.

Tipping, E., J. a. C. Davies, P. A. Henrys, G. J. D. Kirk, A. Lilly, U. Dragosits, E. J. Carnell, A. J. Dore, M. A. Sutton, and S. J. Tomlinson. 2017. Long-term increases in soil carbon due to ecosystem fertilization by atmospheric nitrogen deposition demonstrated by regional-scale modelling and observations. Scientific Reports 7:1890.

U.S. Department of Agriculture. 2015. Summary Report: 2012 National Resources Inventory, Natural Resources Conservation Service, Washington, DC, and Center for Survey Statistics and Methodology, Iowa State University, Ames, Iowa.

Yilmaz, P., L. W. Parfrey, P. Yarza, J. Gerken, E. Pruesse, C. Quast, T. Schweer, J. Peplies, W. Ludwig, and F. O. Glöckner. 2014. The SILVA and “All-species Living Tree Project (LTP)” taxonomic frameworks. Nucleic Acids Research 42:D643–648.

